# Waveform-dependent mechanical signatures shape human vibrotactile perception

**DOI:** 10.1101/2025.11.10.687548

**Authors:** Sarah Bonnet, Roger Holmes Watkins, Rebecca Ericsson, Heidy Daumas, Rochelle Ackerley

## Abstract

Understanding how mechanical vibrations applied to the skin translate into touch sensations is key to advancing tactile interfaces, from non-invasive neuroprosthetics to effective haptic tools. We investigate how modulating vibration waveform, frequency, and amplitude of a wrist-worn vibrotactile device influence perceived touch intensity, while simultaneously measuring accelerations at the actuator and skin surface. Thirty participants provided free-magnitude estimates of perceived intensity, showing that perceived intensity is significantly influenced by all parameters, while being strongly correlated with actuator and skin accelerations. Modulating the waveform shape showed that square waves are consistently rated as more intense. Vibration frequency exerted a non-linear influence, with perceived intensity peaking ∼150 Hz and secondary acceleration and perceptual peaks were found at low frequencies for square and sawtooth waveforms. Thus, the skin faithfully preserves the mechanical signature of vibrations, which are clearly differentiated perceptually. These findings demonstrate that precise tactile characterization, both physically and perceptually, is essential, where device vibrations delivered to the skin determine what is perceived to a very high degree. As vibrations are relatively easy to control and apply, this opens up opportunities to convey a multitude of sensations, from brief taps to pressure, as well as more complex percepts like roughness and texture.

**Summary & Graphical Abstract:** Vibrotactile perception away from the hand remains poorly characterized, yet wearable haptic devices increasingly rely on it for meaningful feedback. Combining perceived intensity ratings with simultaneous acceleration measurements at a vibrotactile wristband and the skin surface, this work shows that perceived intensity is jointly shaped by actuator resonance frequency and harmonic content, with skin-transmitted acceleration emerging as a reliable predictor.

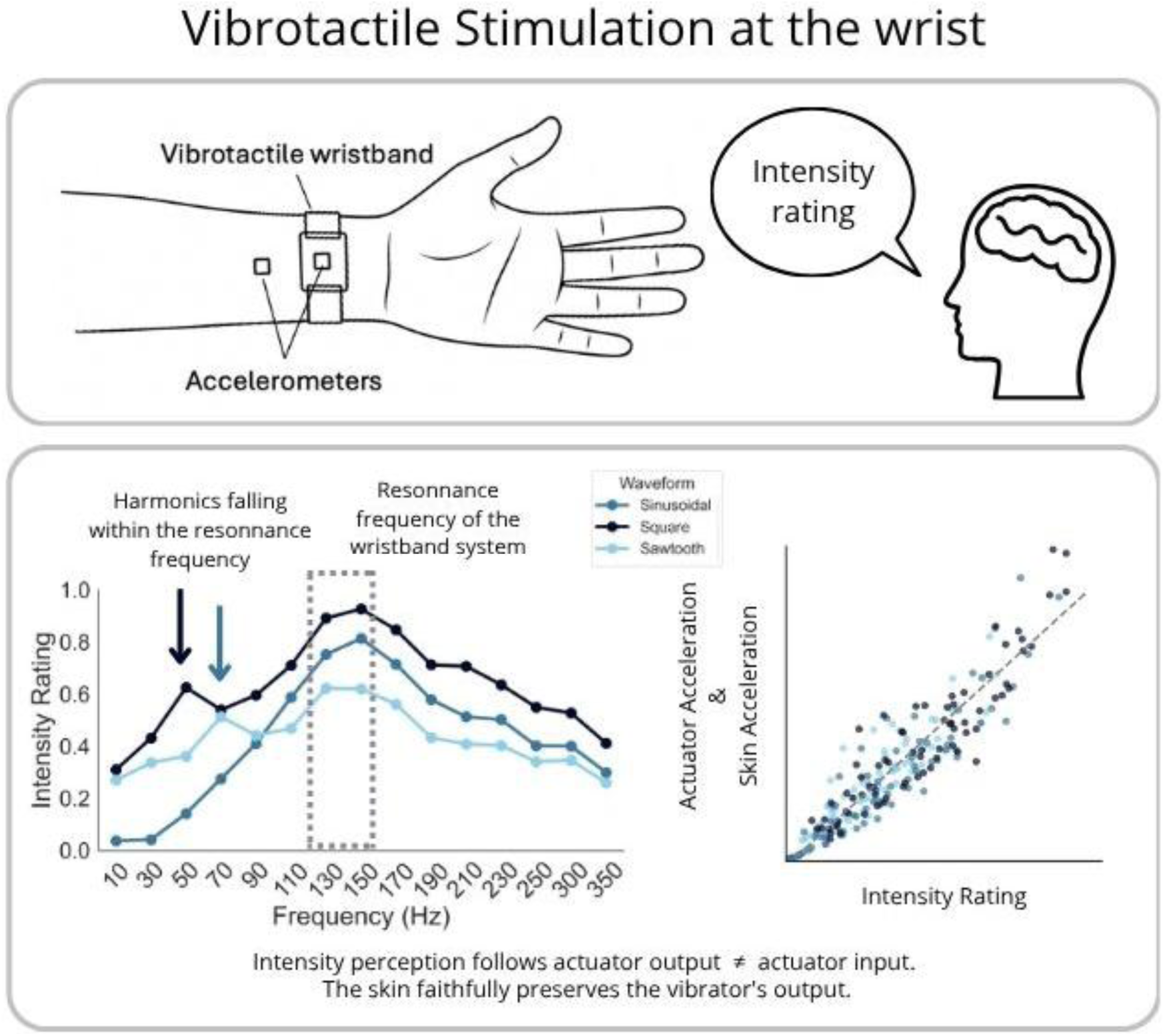

## 1. Introduction

Touch is a fundamental sensory modality that enables us to interact precisely with our environment, control movements, and discern the physical characteristics of objects. Among tactile stimuli, mechanical vibrations applied to the skin play a crucial role for fine texture discrimination, tool manipulation, and haptic communication.^[1], [2], [3]^ Understanding how these vibrations are encoded by the touch system and linked with perception is essential not only for fundamental neuroscience, but also for developing technology, such as sensory substitution devices, neurorehabilitation tools, and precise haptic feedback systems for virtual reality. Extensive psychophysical studies have mapped human tactile detection thresholds and spatial discrimination capacities for vibrotactile stimuli based on their physical parameters. While all mechanoreceptors encode vibration to some extent, they show distinct frequency sensitivities. Pacinian corpuscles, linked to the fast-adapting type 2 (FA-II) mechanoreceptive afferents, are particularly highly sensitive detectors of vibration, responding faithfully to vibrations up to at least 500 Hz and reaching maximal sensitivity in the 200-300 Hz range,^[4], [5], [6], [7]^ making them the principal neural conduit for perceiving fine texture and even remotely transmitted mechanical cues. ^[8], [9], [10]^

Perceptual tactile vibration thresholds on the non-hairy, glabrous skin of the hand form a U-shaped curve across frequencies, with greatest sensitivity around 200-300 Hz,^[11]^ which corresponds well with the maximal sensitivity of Pacinian corpuscles.^[12]^ Perceived intensity generally follows Stevens’ power law, showing a logarithmic-like growth as stimulus amplitude increases.^[13]^ However, in exploring this, studies have predominantly manipulated stimulus amplitude and frequency in isolation, typically employing sinusoidal waveforms under controlled conditions. Despite knowledge of tactile detection thresholds, the mechanisms by which suprathreshold vibrotactile signals translate into graded subjective sensations of intensity remain less understood, with recent studies suggesting that such perceptions depend on the frequency-dependent encoding of afferent activity ^[14]^ and on the dynamic updating of internal decision criteria based on sensory history.^[15]^ Recent evidence suggests a complex interaction between amplitude and frequency: at Pacinian-sensitive frequencies (∼200-300 Hz), minor amplitude changes significantly affected perceived intensity, whereas at lower ‘flutter’ frequencies (< 50 Hz), greater amplitude changes were needed for similar perceptual impacts.^[16], [17]^ Additionally, waveform shape can modulate perceived intensity, with transient- rich signals like square waves feeling more intense than sinusoidal waveforms. ^[18], [19], [20]^ In a recent a study, participants rated waveforms in order of perceived intensity, where stimuli were presented with the same frequency and duration, but all had different waveforms. Square waves were rated as the most intense, followed by sawtooth, and the sinusoidal waveform was rated the least intense.^[18]^ One possible explanation is that square waves generate more high-frequency components that more effectively stimulate the Pacinian channel, which are very sensitive to such high-frequency, transient vibrations.

The question remains about how such vibrations travel through the skin and underlying tissues and how this is linked to perception. Human skin is comprised of several viscoelastic layers of tissue, deformed by mechanical stimulation depending on elastic modulus, viscosity, layer thickness, and boundary conditions, like proximity to muscle, connective tissue, or bone. Evidence has shown that the skin exhibits distinct resonant frequencies, typically ranging from 200-250 Hz. It has been suggested that the viscoelastic properties of the skin and underlying tissues slows the propagation of mechanical waves as the frequency increases.^[21], [22]^ When vibrations are above 100 Hz, the skin has a dampening effect on mechanical vibrations in the skin. ^[23], [24]^ Propagation is also profoundly shaped by the nature of the contact event. Surface waves generated by tapping differ in spectral content and spatial spread from those evoked by gripping or sliding, and the waveform remains surprisingly well preserved as it travels.^[22], [25]^ Greater finger force, or the simultaneous use of multiple digits, boosts vibration intensity and extends its reach.^[25]^ As these biomechanical interactions are non-linear, the skin functions not merely as a passive conduit, but as a frequency-selective filter, which relays information in the optimal Pacinian afferent response range.^[22], [26]^

When a vibratory stimulus contacts the skin, its mechanical energy spreads through surface waves, recruiting mechanoreceptors beyond the initial locus of stimulation.^[26], [27]^ Studies have shown that waves from touch to the finger can traverse the entire palm, cross the wrist, and possibly travel farther along the arm.^[25], [28]^ Remarkably, people who lack tactile input in their fingertips, either congenitally or due to transient local anesthesia, can still judge surface roughness by relying on the residual vibrations that reach more proximal skin regions, such as the wrist.^[29]^ Most research on vibration detection and transmission has focused on fingertip biomechanics, but the anatomical location significantly alters the biomechanical filtering properties. Wearable haptics often aim to target the fingertip, as it is a prime location for exploratory touch; however, haptic devices on the wrist could deliver cues through hairy skin, while remaining close to the glabrous skin. It has even been found that vibration detection thresholds on the volar (inner) wrist are typically higher than the finger.^[30], [31]^ Vibrations applied to the volar side of the wrist, characterized by thicker skin and deeper soft tissue layers, may demonstrate lowered resonance frequencies, increased damping at higher frequencies, and altered perceptual thresholds compared to fingertip stimulation.^[32]^ Consequently, intensity models derived from fingertip data cannot be assumed to generalize to other skin areas. Thus, understanding vibrotactile coding in hairy skin is a pre-requisite for reliable wrist-worn interfaces.

Given the skin’s filtering role, an accurate understanding of tactile perception requires not only knowing the input signal at the actuator, but also quantifying the mechanical signal produced in the stimulated tissues. Measuring this transmitted signal provides a more direct estimate of the stimulation received by mechanoreceptors, and thus of the neural drive underlying perception. In the present study, we aimed to identify which physical properties of a wrist-worn vibratory stimulus most strongly determined perceived intensity, hypothesizing that intensity increases with vibration acceleration and peaks in the 200-300 Hz Pacinian-optimal frequency band. Based on previous literature we expected this to be greatest for square and sawtooth waveforms, and we aimed to perform a detailed examination of how different vibration parameters impact transmission to the skin.

## 2. Results

### 2.1. Differences in intensity to different vibrotactile stimuli

The intensity ratings of the vibrotactile stimuli over the different conditions from 30 participants revealed significant effects of vibrotactile frequency (Friedman ANOVA χ² = 375.02, p < 0.001), amplitude (χ² = 120.00, p < 0.001), and waveform (χ² = 48.27, p < 0.001), although no significant interactions were found between the conditions. For vibration frequency, the perceived intensity showed an inverted-U shape relationship, where there was a rapid increase up to a peak at around 150 Hz that then decreased (**Figure 1A**). Comparisons between frequencies showed significant effects, where a stepwise comparison showed consecutive significant differences from one frequency to the next (stepwise Dunn’s tests all p < 0.001), apart from the step between 130 and 150 Hz (p = 0.171), which represented the peak in the response curve. This effect can also be seen over the different waveforms in the Generalized Additive Mixed Model (GAMM) in **Figure 2** and in the perceived intensity per waveform shape in **Figure 3C**.

**Figure 1.**
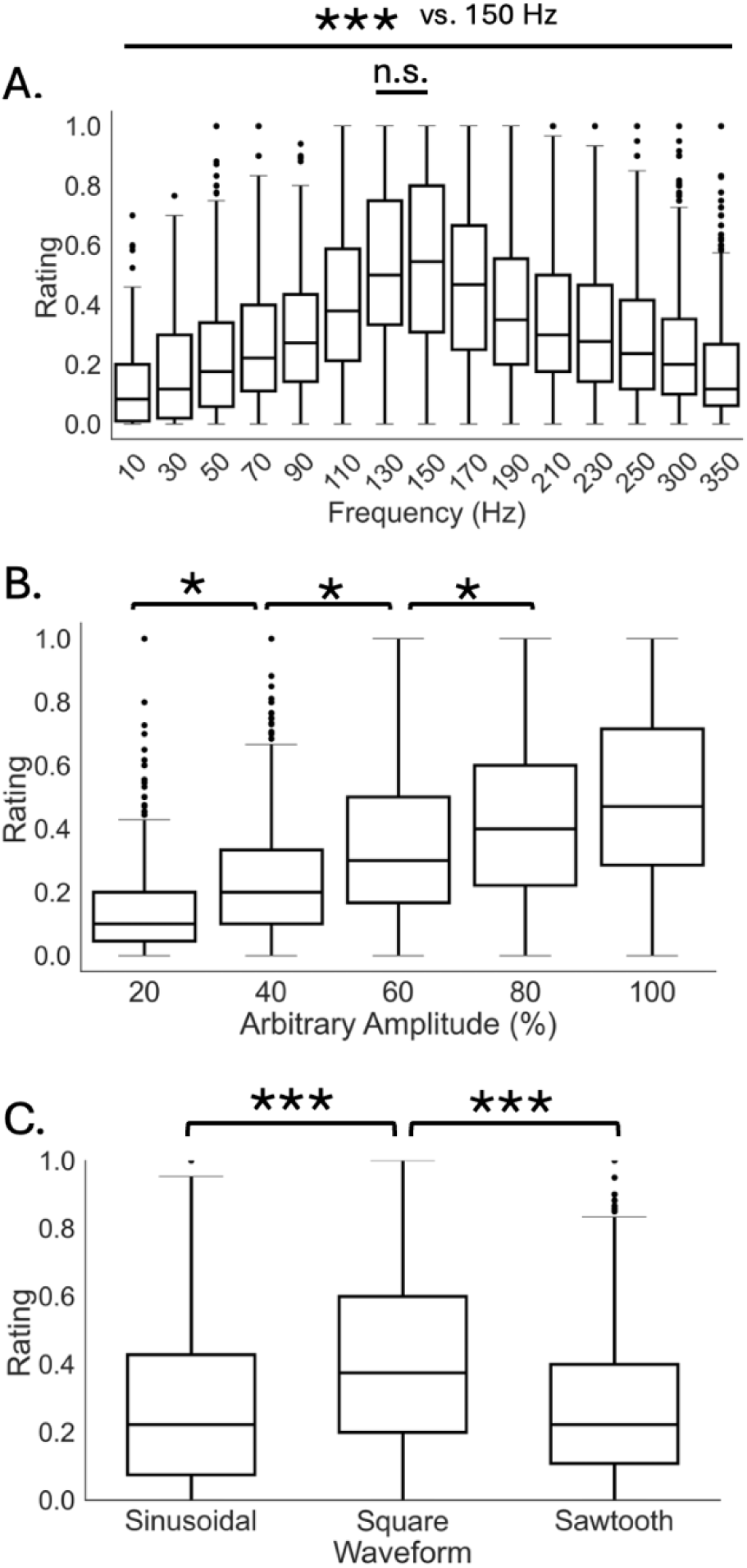
Effects of stimulus parameters on perceived vibration intensity ratings. A. Vibration intensity rating over all waveforms combined at each frequency, where stepwise comparisons from one frequency to the next all showed significant differences (***p < 0.001), apart from between 130 and 150 Hz, which was not significantly different (p = 0.171). B. Vibration intensity rating over all waveforms for each amplitude, where stepwise comparisons from one amplitude to the next all showed significant differences (*p < 0.05). C. Vibration intensity rating for each frequency where all comparisons between all waveform shapes were significantly different (***p < 0.001).

**Figure 2.**
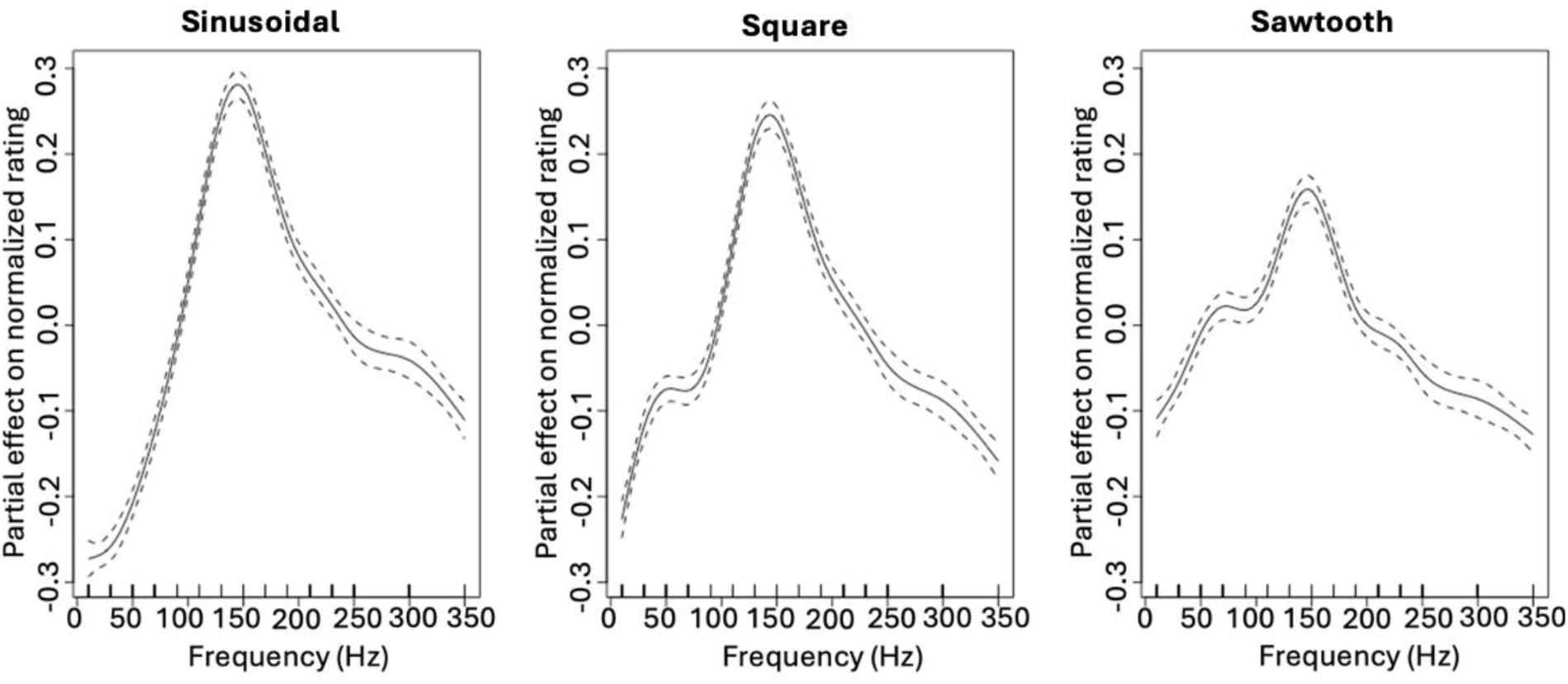
Smooth effects of frequency on normalized ratings for different waveform shapes across the three Generalized Additive Mixed Models (GAMMs). Partial effects of frequency (Hz) on normalized perceptual ratings are shown for sinusoidal, square, and sawtooth waveforms. Dashed lines represent 95% confidence intervals around the estimated smooth function.

**Figure 3.**
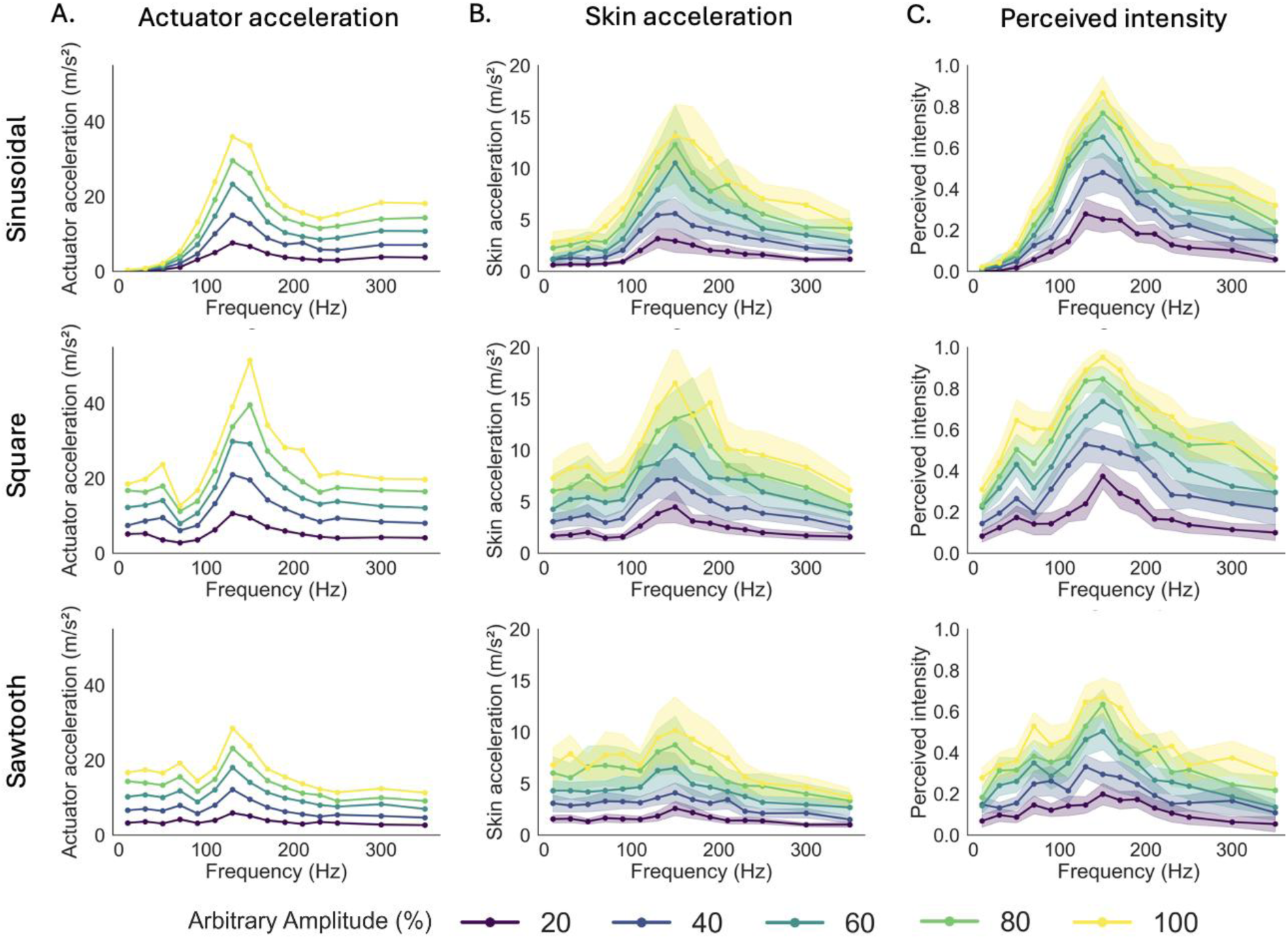
Relationships between perceived intensity, actuator acceleration, and skin acceleration across frequencies and waveform types. A. Actuator peak-to-peak acceleration (m/s^2^) recorded for each frequency for three waveform types: sinusoidal (top), square (middle), and sawtooth (bottom). B. Skin peak-to-peak acceleration (m/s^2^) recorded on the participant’s skin surface at 1 cm from the actuator under the same conditions. C. Perceived intensity of vibrations as a function of stimulation frequency for three waveforms. In all panels, colored lines represent different amplitude levels (20, 40, 60, 80, 100%), as indicated in the legend. Shaded regions denote represent 95% confidence intervals around the mean.

A monotonic increase in perceived intensity was found for vibration amplitude for all waveforms and frequencies, as can be seen in **Figure 1B**, where each step from 20 to 40, 40 to 60, and 60 to 80 (arbitrary amplitude in %) was significantly different (stepwise Dunn’s tests all p < 0.05). These effects can also be seen in **Figure 3C** in the increasing-colored lines at each amplitude over each waveform shape condition.

For the waveform shape, square waves were rated as significantly more intense than sinusoidal and sawtooth waveforms, as can be seen in **Figure 1C** (Dunn’s tests both p < 0.001, no significant difference between sinusoidal and sawtooth waveforms). **Figure 2 and 3C** also display these differences in the model and perceived ratings, respectively.

We tested linear regression models to explore the prediction of intensity from the different conditions, but these yielded low adjusted R² values and failed to capture the non-linear patterns present in the data. Thus, we opted for a GAMM to overcome these issues. The GAMM included the predictor variables of vibratory frequency, waveform shape, and amplitude. The model gave an adjusted R² of 0.59 (scale estimate = 0.020), where both waveform shape and amplitude influenced perceived intensity (**Figure 2**). The sawtooth waveform elicited significantly lower intensity ratings, as compared to both sinusoidal (estimate = 0.014, p = 0.001) and square (estimate = 0.149, p < 0.001) waveforms. Amplitude had a strong, monotonic effect, with perceived intensity increasing progressively from 40% amplitude (estimate = 0.104, p < 0.001) to 100% amplitude (estimate = 0.359, p < 0.001). Frequency showed significant non-linear effects across all waveforms (sinusoidal: F = 380.67, p < 0.001; square: F = 231.15, p < 0.001; sawtooth: F = 87.39, p < 0.001), indicating complex shape-specific modulation of perceptual ratings. Overall, the GAMM offered an accurate representation of the different frequency peaks, especially capturing the square and sawtooth waveforms perceived intensity patterns.

### 2.2. Relationships between perceived intensity and acceleration measures

We wanted to investigate the relationships between perceived intensity ratings and accelerometer outputs, from both the actuator and skin. **Figure 3** shows plots of these measures across all conditions, where it is clear that the perceived intensity and accelerometer measures are highly related, with the forms all showing similar patterns. As per the perceptual results shown in **Figure 1** and **2**, the non-linear effects of frequency can be seen for all measures, where frequency demonstrated an inverted-U shape relationship that peaked around 150 Hz, with secondary lower peaks over all measures for the square and sawtooth waveforms. Further, all measures show monotonic increases in amplitude and general differences between waveforms, where values were higher for square waveforms and lower for sawtooth. Additionally, the correspondence between the two accelerometers can be seen (**Figure 3A** and **3B**), but the skin accelerometer readings were consistently around half that found at the actuator.

To quantify these relationships, we performed correlations between the measures. These gave highly significant positive relationships, namely between perceived intensity and the actuator accelerations (R = 0.932, n = 225, Fisher’s z = 1.62, slope = 0.023 p < 0.001; **Figure 4A**), between the perceived intensity and the skin accelerations (R = 0.950, n = 225, Fisher’s z = 1.84, slope = 0.063 p < 0.001; **Figure 4B**), and between the actuator and skin accelerations (R = 0.946, n = 225, Fisher’s z = 1.79, slope = 0.354, p < 0.001; **Figure 4C**).

**Figure 4.**
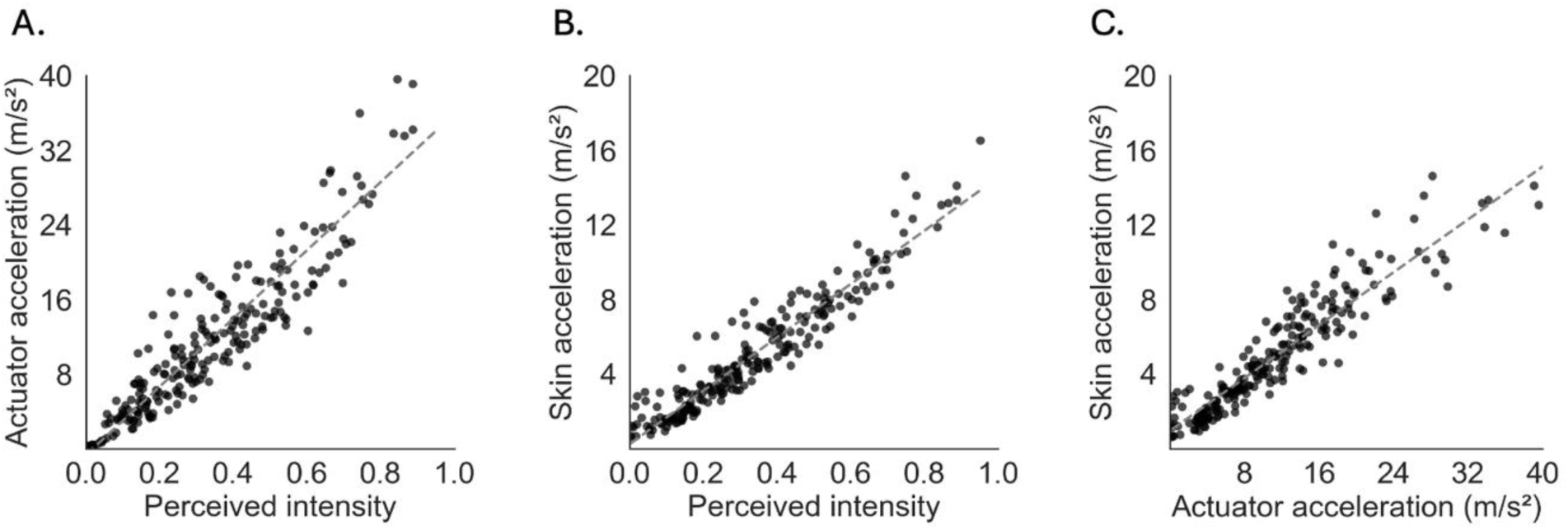
Correlations between perceived intensity and the accelerometer outputs. Scatter plots illustrate the relationships between: A. perceived intensity ratings and actuator acceleration (m/s²); B. Perceived intensity ratings and skin acceleration (m/s²); C. Actuator and skin acceleration (m/s²). Spearman’s rank correlations revealed strong positive associations across all comparisons. Data from 20 participants over a total of 225 conditions.

### 2.3. Spectral analysis of actuator and skin vibration signals

To further characterize the mechanical output of the actuator and its transmission to the skin, we performed a spectral analysis of the accelerometer signals across all controlled frequencies and waveform shapes (**Figure 5**). Across all three waveforms, a consistent band of elevated spectral power was observed around 130-150 Hz in both the actuator and skin signals, reflecting the resonance frequency of the actuator-wristband-skin system (**Figure 5A and 5B**). For the sinusoidal waveforms, the dominant spectral component closely tracked the commanded frequency across the tested range, visible as a diagonal ridge in the actuator spectral power plots (**Figure 5A**, top). Below 30 Hz, however, no clear spectral peak was observed at the commanded frequency; instead, the actuator produced diffuse, low-power output spread across a broad frequency range. This is consistent with a failure of the device to generate coherent vibration at these frequencies. For square and sawtooth waveforms, this resonance band was further amplified by harmonic components falling within this range. At a commanded frequency of 50 Hz for the square waveform, the dominant spectral component was the third harmonic at ∼150 Hz, falling directly within the resonant band. Similarly, for the sawtooth waveform at 70 Hz, the second harmonic at ∼140 Hz carried the greatest spectral power. In both cases, the device was effectively vibrating at its resonant frequency regardless of the commanded fundamental, providing a direct mechanical explanation for the secondary perceptual intensity peaks observed at these frequencies (**Figure 3C**).

**Figure 5.**
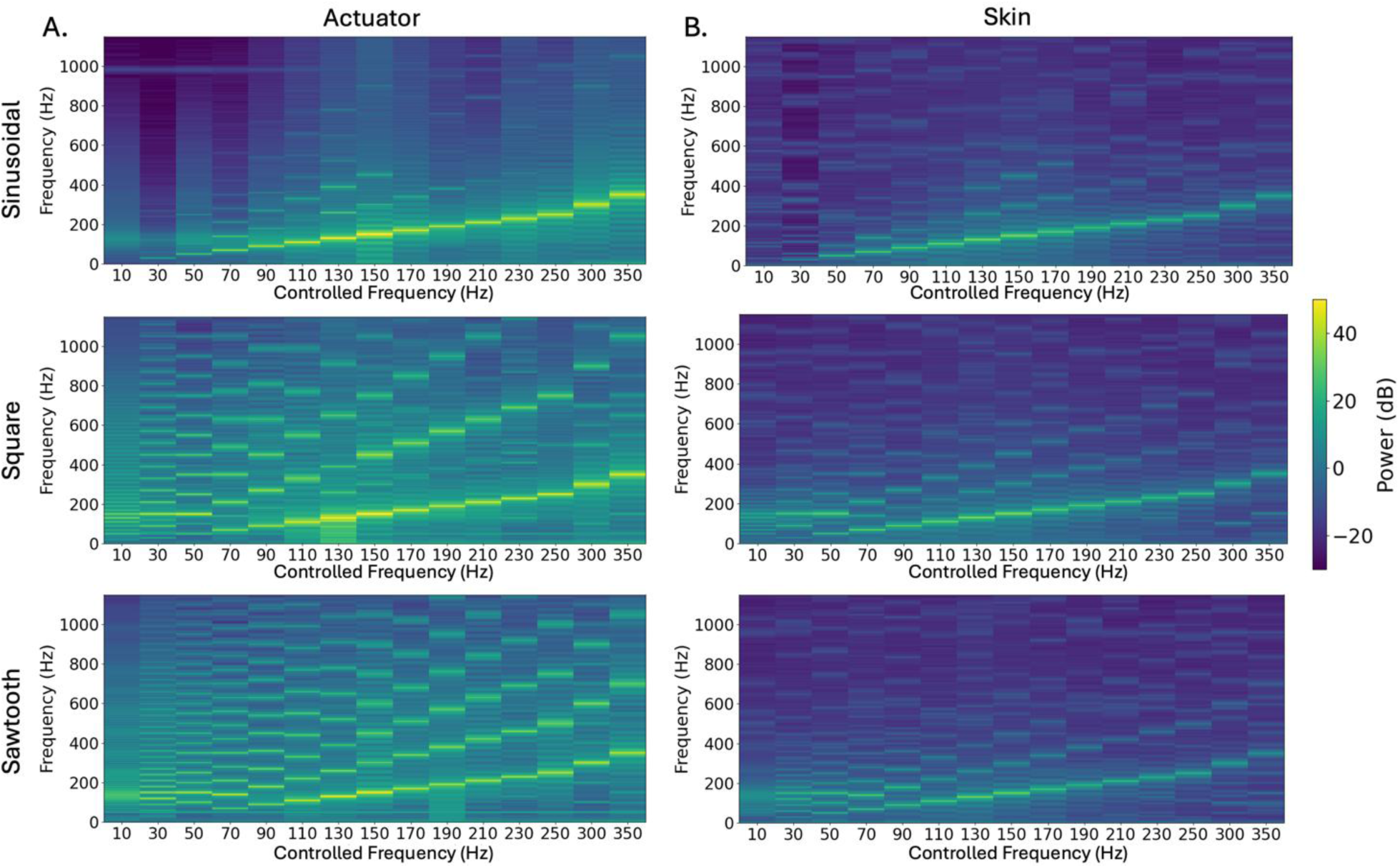
Spectral power of accelerometer signals at the actuator and skin surface across controlled frequencies and waveform shapes. Power spectral density (dB) is shown as a function of recorded frequency (y-axis) and controlled frequency (x-axis), for the actuator (A) and skin (B) accelerometers, across sinusoidal (top), square (middle), and sawtooth (bottom) waveforms. For the sinusoidal waveform, the spectral peak tracked the commanded frequency along the diagonal, except below 30 Hz where no clear peak was observed. Across all waveforms, a consistent increase in spectral power was observed around 130-150 Hz, reflecting the resonance frequency of the actuator-wristband-skin system. For square and sawtooth waveforms, this resonance was further amplified by harmonic components falling within this range: at 50 Hz commanded frequency for the square waveform, the dominant component appeared at ∼150 Hz (third harmonic), and at 70 Hz for the sawtooth, at ∼140 Hz (second harmonic). These patterns were preserved in the skin signals with an overall reduction in power and harmonics.

The skin accelerometer signals (**Figure 5B**) preserved the overall spectral structure observed at the actuator, including the diagonal tracking for sinusoidal stimuli, the resonance amplification at 130-150 Hz, and the harmonic-driven peaks for square and sawtooth waveforms. An overall reduction in spectral power was observed across all conditions, consistent with the attenuation of vibration during propagation through skin tissue.

## 3. Discussion

The present study investigated the physical determinants of perceived vibration intensity using a wrist-worn vibrotactile device, while simultaneously measuring the mechanical signal at both the actuator and the skin surface. We found three principal findings. First, perceived intensity peaked around 150 Hz, not in the 200-300 Hz range predicted by Pacinian corpuscle tuning, but at the resonance frequency of the actuator-wristband-skin system, thus showing how the percept intensity was mainly driven by the physical vibration, rather than being nearer the sensitivity optimum of specific mechanoreceptor types. Second, this resonance-driven peak was preserved in the skin acceleration signal and propagated highly faithfully 1 cm away from the actuator, with the overall profile of the stimulus remaining intact despite an approximate halving of amplitude. Third, secondary perceptual peaks at 50 Hz for square and 70 Hz for sawtooth waveforms were not driven by the commanded fundamental frequency, but by harmonics that fell directly within the device’s resonant range. Taken together, the strong fidelity between actuator behavior, skin acceleration, and perceived intensity demonstrate that the skin is a highly sensitive transducer of actuator dynamics, faithfully encoding not only the intended stimulus parameters, but also device-specific artifacts such as resonance amplification and low-frequency failure, where tactile perception was very sensitive to all these factors, even when focused on the hairy skin of the wrist.

### 3.1. The resonance of the system shapes frequency-dependent perception

A notable finding is that perceived intensity peaked at ∼150 Hz, closer to the resonance frequency of the actuator (∼133 Hz) than to the Pacinian optimum of 200-300 Hz predicted by fingertip psychophysics.^[4], [5], [33], [34], [35], [36]^ The spectral data offer a plausible explanation: across all waveforms, power spectral density showed consistent amplification at 130-150 Hz, reflecting the resonance of the coupled actuator-wristband-skin system, and this amplification was preserved in the skin acceleration signal. However, an important methodological caveat must be acknowledged. In the present study, amplitude was expressed as a percentage of the actuator’s maximum output rather than as a calibrated displacement or force held constant across frequencies. As a consequence, the amplitude of displacement of the actuator, was not equated across frequencies at the time of stimulation. It is therefore possible that the peak in perceived intensity around 150 Hz reflects, at least in part, greater mechanical output at the actuator’s resonant frequency for a given arbitrary amplitude command, rather than a purely perceptual sensitivity for this frequency range. With this caveat in mind, the convergence between the perceptual peak, the actuator resonance, and the spectral amplification observed in both accelerometer signals remains a coherent and informative pattern. Understanding how these dynamically change would therefore allow for the transformation of drive signals to be used for more faithful tactile signal reproduction.

### 3.2. Harmonics as perceptual amplifiers

A particularly compelling aspect of our results concerns the secondary intensity peaks observed for square and sawtooth waveforms at low commanded frequencies. At 50 Hz for the square waveform, the dominant spectral component reaching the skin was not the fundamental, but the third harmonic at ∼150 Hz, within the device’s resonant range. Similarly, at 70 Hz for the sawtooth waveform, the second harmonic at ∼140 Hz carried the most mechanical power. In both cases, participants perceived these stimuli as more intense than their commanded frequency would suggest, because the device was effectively vibrating at its resonant frequency regardless of the nominal input. This finding reframes how waveform shape should be understood in haptic and neuromodulatory design: the perceptual consequences of a waveform are not simply a function of its time-domain appearance, but of how its harmonic content interacts with the resonance profile of the full actuator-wristband-skin system. Designing waveforms that strategically place harmonics within the device’s resonant band may therefore be a more principled approach to intensity modulation than selecting waveform shape based on prior literature alone.^[18]^

Very weak perceived tactile intensity to sinusoidal stimulation at low frequencies (< 30 Hz) were accompanied by a near-absent acceleration signal in both the actuator vibration amplitude and the skin, indicating that the mechanical properties of the actuator did not effectively deliver stimulation down at these frequencies; however, the touch was nerveless sensed. For square and sawtooth waveforms, the spectral picture differed: even at low commanded frequencies, the actuator generated a broad band of energy around its resonance frequency (∼130-150 Hz), which likely explains why perceptual responses, while reduced, were not entirely absent. This waveform-dependent divergence in low-frequency behavior further illustrates that perception at any given commanded frequency cannot be understood without characterizing the stimulation actually delivered by the actuator. It further shows that the waveform can be modulated independently at a fixed frequency to provide a qualitatively-different and perceivable sensation.

### 3.3. Amplitude and waveform shape as independent modulators

Beyond frequency, amplitude emerged as a strong, monotonic determinant of perceived intensity, consistent with prior psychophysical literature demonstrating power-law scaling of tactile perception with stimulus strength. ^[13], [14], [15], [16]^ The most striking finding was that both actuator and skin accelerations proved to be the best predictors of perceived intensity. While acceleration has been shown to be a secondary dimension in vibrotactile discrimination,^[37]^ recent work confirms its value as a robust correlate of perceived intensity,^[38]^ and our results extend this, supporting acceleration as a reliable energetic proxy for the mechanical energy reaching mechanoreceptors, rather than a direct perceptual code.^[32]^ Further, tactile acceleration signals may play a fundamental role in activating mechanoreceptors, especially the vibration-sensitive Pacinian FA-IIs. Temporally-accurate Aβ low-threshold mechanoreceptors are change detectors and fire especially well to rapid mechanical forces, acting effectively like accelerometers. The consistency of the amplitude effect further suggests that it operates as a gain control independent of the spectral shaping introduced by frequency and waveform.

Waveform shape acted as a categorical modulator, with square waves eliciting consistently higher intensity ratings than both sawtooth and sinusoidal waveforms across all frequencies and amplitudes. This aligns well with prior findings^[18], [19], [20]^ and can be attributed to two complementary mechanisms. The sharp transitions of square waves introduce a broad harmonic spectrum that more effectively stimulates Pacinian corpuscles, which are highly sensitive to high-frequency transient content.^[39]^ Although sawtooth waves contain more harmonics than square waves, what determines perceptual intensity is not the total number of harmonics, but rather the concentration of energy in those that fall within the actuator’s resonant band. Square waves contain only odd harmonics (3f, 5f, 7f…) with energy decreasing as 1/n, which concentrates a disproportionate amount of power in the first odd harmonic (3f), which at a commanded frequency of 50 Hz falls at exactly 150 Hz, directly within the resonant range. Sawtooth waves, while richer in harmonic content overall, distribute their energy across both odd and even harmonics (2f, 3f, 4f…), reducing the power of any individual harmonic and making favorable alignment with the resonant band less systematic. This harmonic energy distribution, rather than harmonic richness per se, likely explains why square waves were consistently perceived as most intense despite sawtooth waves being spectrally richer.

### 3.4. Skin transmission preserves the stimulus profile

A key methodological contribution of this study is the direct measurement of vibration transmission from the actuator to the skin surface. The strong correlation between actuator and skin accelerations demonstrates that the overall profile of the mechanical stimulus was faithfully preserved 1 cm away from the actuator, despite an approximately twofold reduction in amplitude. Importantly, this tight coupling suggests that actuator vibration, considerably easier to measure in practice, can serve as a reliable proxy for skin vibration, simplifying the characterization of the actual stimulus delivered to the tissue. This has important implications beyond basic neuroscience. In the context of neuromodulation and neuroprosthetics, where vibrotactile actuators are increasingly used to convey sensory feedback or elicit afferent nerve activity, knowing that device-level measurements faithfully reflect skin-level mechanics is critical for stimulus reproducibility and dose control. It further suggests that skin acceleration could serve as a physiologically grounded target metric for close-loop control of perceived intensity, offering a more principled alternative to actuator-referenced output measures and bringing haptic stimulation closer to the standards of rigor expected in clinical neurostimulation.

### 3.5. Limitations and future directions

Several limitations of the present work should be acknowledged. Most importantly, vibration amplitude was expressed as a percentage of the actuator’s maximum output and was not calibrated to equate displacement or force across frequencies. This means that the mechanical output of the device varied across the frequency range in ways that were not fully controlled, which limits causal interpretation of the frequency-dependent intensity effects. Future studies could employ laser Doppler vibrometry or impedance-matched stimulation to deliver equated displacements across frequencies, allowing a cleaner separation of device mechanics and perceptual sensitivity. However, such systems are costly, technically demanding, and poorly suited for deployment in real-world or wearable devices. In contrast, the accelerometer-based approach used here, while imperfect, is lightweight, low-cost, and readily integrated into practical haptic systems. Our conclusions are also specific to a single linear resonant actuator with a resonance at 133 Hz; whether the patterns observed here generalize to actuators with different resonance frequencies, or to other motor technologies such as eccentric rotating mass motors or piezoelectric transducers, remains to be established. All measurements were obtained with the forearm at rest, whereas everyday use of wrist-worn devices involves varying degrees of muscle tension that stiffen underlying tissues and may shift both mechanical filtering and perceptual thresholds. Finally, our sample comprised young adults; given known age-related changes in skin hydration, tissue stiffness, and mechanoreceptor density,^[40]^ the present intensity model may not generalize directly to older populations, for whom wearable haptic devices are increasingly relevant.

A natural extension of this work would be to combine our protocol with measurement of the evoked mechanoreceptive afferent discharge. This can be achieved using microneurographic recordings, where the activity of single mechanoreceptive afferents to different stimuli can be recorded directly in human subjects, allowing a quantification of the sensory signals generated by a particular stimulus.^[41], [42]^ These FA-II afferents, whose Pacinian corpuscle end-organs are the primary neural substrate of high-frequency vibration perception. Such recordings would clarify whether the perceptual intensity peak around 150 Hz reflects a genuine shift in afferent firing rates driven by the actuator-wristband-skin resonance. A particularly compelling question concerns remote afferent activation: given that surface waves propagate well beyond the contact zone, FA-II afferents located at a distance from the actuator may contribute substantially to perceived intensity. Expanding the accelerometer array along the forearm would allow the spatial profile of vibration propagation to be mapped simultaneously with neural responses, providing a direct mechanistic link between skin mechanics, afferent discharge, and perception.

### 3.6. Implications

Our findings converge on several practical principles at the intersection of somatosensory psychophysics and haptic engineering. Our overarching message is straightforward: in wearable haptics, the vibration command is not the vibration the skin receives, and yet the skin tracks the mechanical fingerprint of the device with remarkable fidelity, along with perception. This has immediate consequences for both sides of the touch science and engineering pipeline. For fundamental research, it means that perceptual effects attributed to the nervous system must first be screened for device-level origins; actuator characterization should be considered as standard a step as participant screening. For engineering, it means that the resonance of the coupled actuator-skin system, the harmonic structure of the waveform, and amplitude modulation together provide a coherent, relatively orthogonal toolkit for designing graded haptic sensations, one grounded in measured skin mechanics rather than in fingertip psychophysics alone. As haptic interfaces move toward richer, more nuanced feedback in human-machine interaction and neural engineering, bridging the gap between what a device is commanded to do and what the skin actually receives will be essential to delivering reliable, calibrated, and perceptually meaningful tactile experiences.

## 4. Methods

### Participants

Thirty healthy volunteers participated in the full experiment (15 females and 26 right-handed by self-report; age range: 18-32 years, mean age: 25 years). In 10 of these participants, we only gained intensity ratings. The study was approved by an ethics committee (Comité de Protection des Personnes Ouest III, approval number: 23.03773.000242) and conformed to the Declaration of Helsinki, apart from pre-registration. All participants were given standard information about the experiment, signed an informed consent form, and received financial compensation for their time. Inclusion criteria required participants to be at least 18 years of age, with no restriction based on handedness. Exclusion criteria included conditions potentially affecting tactile sensitivity, such as neurological or psychiatric disorders, epilepsy, diabetic neuropathy, dermatological conditions, pregnancy or breastfeeding, and being under legal protection.

### Experimental setup

Participants first completed a demographic questionnaire (age, gender, handedness, occupation). Skin temperature was measured at the wrist and index finger pad using an infrared thermometer (AP-1580 AOPUTTRIVER), as well as skin water content (Skin Analyser, Hurrise)^[43]^ as control measures, both before and after the experiment. These demographic and skin measures were taken as controls, but were found to have no significant effects on the results. Ambient room temperature was also recorded at the start of the session (mean: 25.1 ±1.3°C). All tests were conducted on the side of the participant’s dominant hand.

Participants were seated comfortably on an adjustable chair, with the arm of their dominant hand positioned palm-side up facing away from them, supported by a cushion, so that they could rest in a comfortable position. They were then fitted with a custom 3D-printed vibrotactile wristband (Haptify) housing a voice-coil actuator with a resonance frequency at 133 ±13 Hz, which had an accelerometer (ADXL326, ±19 G, Analog Devices) embedded between the actuator and the inner thermoplastic polyurethane housing (hardness 95A). The wristband was positioned on the wrist of their dominant hand, ∼ 2 cm proximal to the wrist crease, and aligned with the flexor tendons (**Figure 6A**). The high-range accelerometer in the wristband was PCB mounted (Adafruit ADXL326 breakout, product no. 1018) and connected, to a data acquisition system (National Instruments USB-6216). To quantify vibration propagation through the skin, a different, miniaturized, high-sensitivity accelerometer (LIS344ALH, ±2 G, STMicroelectronics) was mounted on a custom lightweight printed circuit board including an onboard power capacitor and an LED for positional tracking. The device was connected to an output amplifier circuit via a low resistance and capacitance 70mm flat flexible cable (Molex Premoflex) along with 1 nF band limiting capacitors to achieve a bandwidth >1 kHz with minimal mechanical filtering imposed by the connectors.

**Figure 6.**
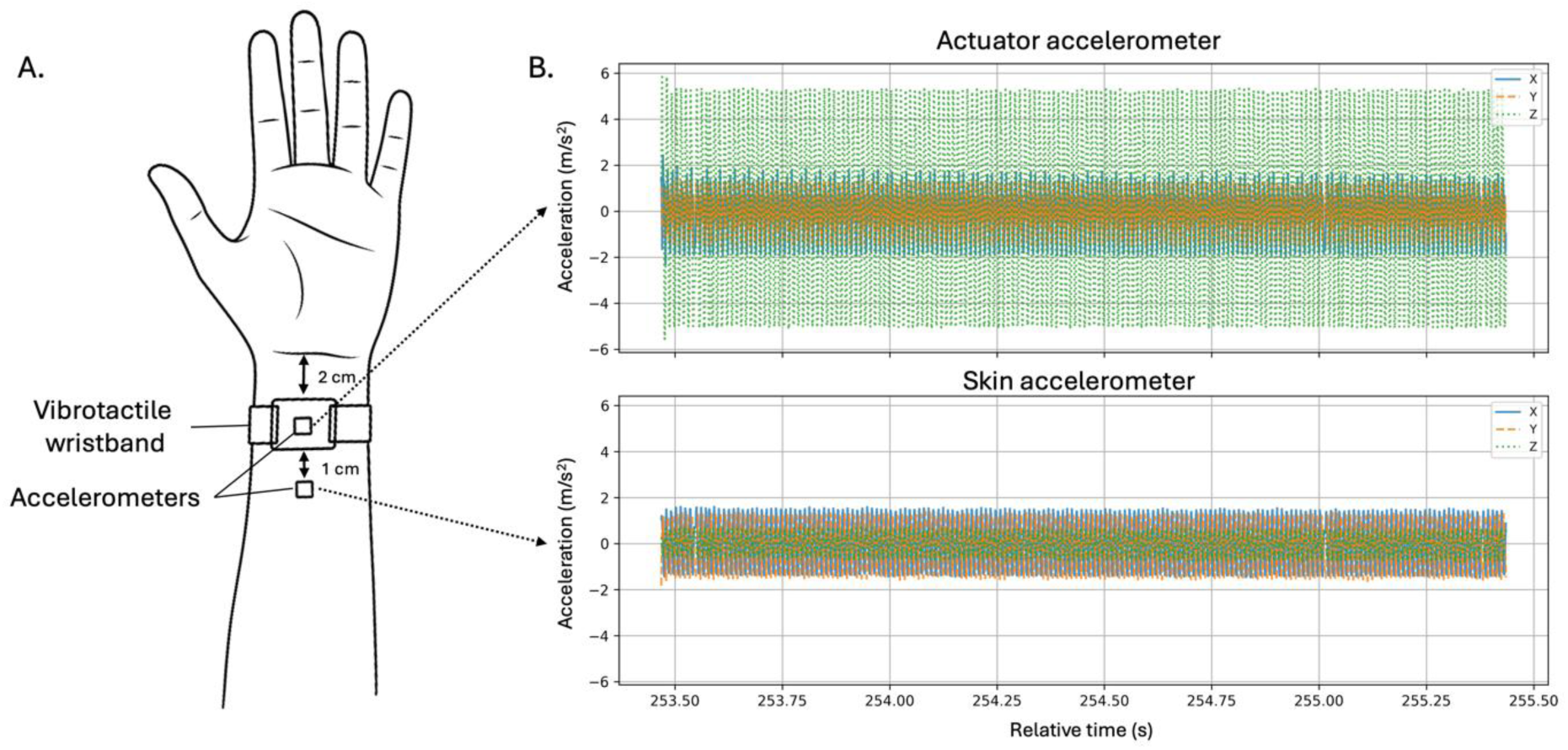
Schematic illustration of the vibrotactile wristband setup and baseline accelerations from the wristband. A. The 3D-printed wristband, incorporating an embedded accelerometer, was positioned 2 cm proximal to the wrist crease on the dominant hand. The wristband was adjusted for each participant to give consistent skin contact for reliable experimental delivery. A second accelerometer was placed 1 cm proximal to the wristband on the inner forearm to measure vibration propagation through the skin. B. Examples of recorded accelerations at the actuator and in the skin.

The amplifier board scaled the accelerometer signals and output ±3.3 V for high-fidelity transmission of acceleration signals over long cables to the data acquisition system (NI USB- 6216). This configuration enabled real-time acquisition of the full acceleration range for each site, where the vibration at the wristband was much higher than in the skin. A gravity-based calibration was performed to validate the accelerometer outputs in different orientations. Custom Python software was developed to record and visualize tri-axial acceleration data from both sensor types. Using the *nidaqmx* library, signals were sampled at 3 kHz per channel via the NI USB-6216. Real-time visualization was implemented with *DearPyGui*, and time-stamped data were logged in CSV format for offline analysis.

The system utilized a digitally controllable Class-D amplifier (MAX 98357) connected to a WiFi/Bluetooth capable microcontroller to enable real-time control of vibration patterns. The microcontroller received command signals from the control software via USB to control the actuator waveform shape, frequency, and amplitude. Custom firmware written in C++ (implemented in the Arduino IDE) allowed the microcontroller to receive commands from the experimental computer, parse messages, and stream signals to the amplifier at 8 kHz sampling rate and 16-bit resolution. This setup ensured low-latency, high-fidelity delivery of time-varying tactile stimuli synchronized with experimental events.

### Experimental Task and Parameters

During the experiment, participants kept their eyes closed and wore noise-cancelling headphones to eliminate any audible cues of the vibration characteristics (3M; 30 dB attenuation). They received 225 randomized vibratory stimuli, each lasting 2 s, that were separated by 10 s rest intervals, where they also rated the stimulus intensity. Three parameters, each with different levels, were tested, namely: frequency (15 levels: 10, 30, 50, 70, 90, 110, 130, 150, 170, 190, 210, 230, 250, 300, 350 Hz), amplitude (5 levels: 20, 40, 60, 80, 100 arbitrary units, as %), and waveform shape (3 levels: sinusoidal, square, sawtooth). Parameter combinations were randomized for each participant. After each stimulus, participants rated the perceived intensity verbally as a magnitude estimate on an unbounded numerical scale that overcomes end-point limitation issues.^[44]^ For these free numeric ratings, the participants were instructed that zero indicated "no vibration sensation" and was the lower limit (i.e. no negative numbers were allowed, but the participants could use decimal places). They were asked to provide a starting rating for the vibration, then instructed that for subsequent vibrations, they could use a higher number if the sensation was more intense and a lower number if the sensation was less intense. The scale was open-ended, meaning that there was no upper limit and the participants could choose their own scale range. The verbal rating was noted electronically after each vibration stimulus.

### Control measures and accelerometer analyses

Prior to the experiment, we recorded the vibrator’s peak-to-peak accelerations as a function of all the frequency, amplitude, and waveform conditions. These measured accelerations served as a quantification of the vibrational output of the wristband when attached to the wrist under typical experimental conditions from one participant (**Figure 6B**). These data were used as a baseline, control condition in further analyses and are referred to as the actuator acceleration (m/s^2^).

Raw voltage recordings from both the actuator accelerometer and the skin-mounted accelerometer were offset-corrected and converted into SI units (m/s²) by subtracting each sensor’s reference voltage, multiplying by its sensitivity, and then transforming G-forces into peak-to-peak acceleration (m/s²). After voltage-to-acceleration conversion, the mean of every axis was removed to eliminate residual DC components. For both sensors, we retained the three orthogonal channels (X, Y, and Z) and computed the following Euclidean norm (Equation 1) to capture overall motion:

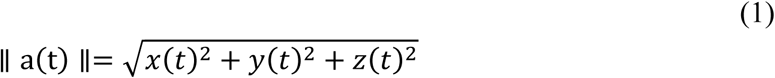

The active vibration window was then identified automatically by applying a median filter to the magnitude signal, defining a threshold at the 10^th^ percentile plus 20% of the signal’s dynamic range, and marking the first and last time points above that threshold. Peak-to-peak acceleration was obtained for each axis (and for the 3-D vector magnitude) by subtracting the minimum from the maximum value of the DC-centered acceleration trace across the entire segment.

To characterize the spectral output of the actuator and its transmission to the skin, power spectral density (PSD) was computed for each frequency, amplitude, and waveform condition using a multitaper time-frequency approach. Raw accelerometer signals were first high-pass filtered (Butterworth, 4th order, cutoff = 1 Hz) and resampled to a uniform time grid. For each recording, a multitaper time-frequency representation was computed on the Z-axis acceleration signal, and the resulting power was averaged over the stable portion of the stimulus window (excluding 0.5 s at each edge to avoid transient effects).

### Statistical analyses

Data were analyzed using Python (version 3.10) within the PyCharm IDE (JetBrains, version 2024.2.2). Individual perceptual intensity ratings were normalized within each participant using a min-max scaling to the [0, 1] range for all subsequent analyses. Exact p values are given from p < 0.05 to p < 0.001. Normality of the distribution of intensity ratings was assessed using Shapiro-Wilk tests, which showed significant deviations from normality across conditions, justifying the use of non-parametric approaches. We used Friedman ANOVA tests to examine significant differences between the perceived intensity of the vibration conditions, and when significant, we performed pairwise post-hoc comparisons using Dunn’s tests corrected for multiple comparisons via false discovery rate correction.

We then conducted analyses to investigate the prediction of perceived intensity from the different vibration parameters. Regression models were tested to best fit the data per condition, but classic regressions with linear, quadratic, and cubic fits left a substantial amount of variance unexplained. We therefore opted for a Generalized Additive Mixed Model (GAMM), implemented in RStudio (version 2024.12.1+563; Posit Software, PBC, Boston, MA, USA). GAMMs allow for flexible modeling of non-linear relationships (e.g. the effect of frequency) through smooth functions while accounting for inter-individual variability using random effects. To investigate relationships between measured data, Spearman’s rank correlations were conducted to assess the relationships between perceived intensity ratings and accelerometer outputs (both from the actuator and skin), with effect sizes expressed as Fisher’s *z*.

## Acknowledgements

The present study was co-funded by Haptify.

## Data Availability Statement

The perceptual data are available in the OSF repository https://osf.io/j487e and the acceleration data are available on request to the authors, due to their large size.

